# Chromatin states at homeoprotein loci distinguish axolotl limb segments prior to regeneration

**DOI:** 10.1101/2022.11.14.516253

**Authors:** Akane Kawaguchi, Jingkui Wang, Dunja Knapp, Prayag Murawala, Sergej Nowoshilow, Wouter Masselink, Yuka Taniguchi-Sugiura, Jifeng Fei, Elly M. Tanaka

**Author notes:** These authors contributed equally to this work (Akane Kawaguchi, Jingkui Wang). National Institute of Genetics, Mishima, Shizuoka, Japan (Akane Kawaguchi). Department of computational biology and digital sciences, Boehringer-Ingelheim RCV, Vienna, Austria.

## Abstract

The salamander limb regenerates only the missing portion. Each limb segment can only form segments equivalent to- or more distal to their own identity, relying on a property termed “positional information”. How positional information is encoded in limb cells has been unknown. By cell-type-specific chromatin profiling of upper arm, lower arm, and hand, we found segment-specific levels of histone H3K27me3 at limb homeoprotein gene loci but not their upstream regulators, constituting an intrinsic segment information code. During regeneration, regeneration-specific regulatory elements became active prior to the re-appearance of developmental regulatory elements. This means that, in the hand segment, the permissive chromatin state of the hand homeoprotein gene *HoxA13* engages with regeneration regulatory elements, bypassing the upper limb program.

## Introduction

Amputation anywhere along the length of the salamander limb yields regeneration of only the missing portion (For review, see (*2*) (*3*) (*4*). In salamanders such as the axolotl, limb fibroblasts and other lateral plate mesoderm descendants (Connective Tissue, CT cells) harbor stable, segment-specific properties, often called positional information (*5*) (*6*) (*7*). Labeled hand CT cells transplanted into the mature upper arm enter the blastema but then contribute only to the regenerated hand. In contrast upper arm CT cells contribute to upper arm, lower arm, and hand (*6*). While cell surface proteins that affect blastema cell sorting, PROD1 and TIG1, may show a shallow gradient of expression in the adult limb (*8*, *9*), how resident limb CT cells are intrinsically and molecularly poised to initiate segment-appropriate regeneration programs has remained unknown.

Vertebrate limb segments have a molecularly-defined origin in the developing limb bud where a series of transcription factors are expressed under the influence of spatio-temporally-modulated signaling (For review see: (*10*) (*11*)). The MEIS homeoproteins, MEIS1,2,3 play a critical role in specifying upper arm cells and suppressing hand identity. In the early limb bud, MEIS expression is found throughout the limb bud but as the limb bud grows beyond proximalizing signals and influenced by distal ectodermal signals such as Fgf and Wnts, a graded MEIS-low region at the distal tip emerges (*12*–*15*). This distal region expresses *HoxA11*,associated with forearm morphogenesis, and then under an apparently autonomous timer, *HoxA13* which is associated with hand formation (*16*, *17*). Global *Meis* expression induced by viral over-expression or exposure to retinoic acid (RA) prevents distalization of progenitors including suppression of *HoxA13* (*12*, *13*, *18*). Finally, limb conditional mouse mutants of *Meis1,2* lack upper limb elements (*15*, *19*). The *Shox2* gene is also implicated in upper limb morphogenesis. This gene is expressed in the proximal bud and its genetic deletion in the mouse limb leads to loss of upper limb structures and has been proposed to be involved in early cartilage differentiation (*20*, *21*). These studies show the functional involvement of MEIS and SHOX homeoproteins in the specific development of upper limb structures.

The progressive expression of 5’*HoxA/D* genes is essential for the elaboration of limb segments, with *Hox13* genes playing a critical role in transitioning to the hand program by suppressing upper limb programs. During limb development, *HoxA9* then *HoxA11* followed by *HoxA13* are expressed along the proximal-distal axis (*22*) (*23*). The genetic disruption of *HoxA13* and its paralog *HoxD13* in mouse shows loss of hand structures (*24*). In the salamander, *Pleurodeles waltl*, disruption of *HoxA13* alone is sufficient to obliterate the formation of hand elements during development (*25*). Over-expression of *HoxA13* throughout the chick limb bud truncates lower wing structures and *in vitro HoxA13*-expressing cells sort away from non-expressing cells (*26*). At the genetic level, HOXA13 binds to gene regulatory elements in the *HoxA11* gene regulatory region and up-regulates a *HoxA11* anti-sense product to terminate *HoxA11* expression in the primordial hand element (*27*). Furthermore, *Hox D13* expression in the chick limb bud suppressed *Meis2* which is also a target of *HoxA13*, showing that *Hox13* expression acts to antagonize the MEIS-driven upper arm program (*12*, *13*, *28*). Taken together these studies established MEIS and HOXA13 as important, counteracting factors that control upper arm versus hand development.

MEIS and HOXA13 proteins also play important roles in axolotl limb regeneration and display level-specific protein induction depending on the site of amputation. Upon upper arm amputation, nuclear MEIS proteins are expressed in CT-derived blastema cells throughout the early blastema while they are not expressed in CT-derived blastema cells after hand amputation (*6*). In contrast, HOXA13 is not expressed in the early upper arm blastema, but is expressed throughout the early hand blastema (*29*), showing that upper arm versus hand CT cells initiate expression of different homeoproteins corresponding to the structures they will regenerate. This hand identity state is an intrinsically stable state persisting through amputation, since labeled, mature CT cells transplanted into the upper arm prior to amputation results in those cells only contributing to the hand (*5*, *6*).

An upper arm blastema must ultimately regenerate the entire limb. At mid-to-late bud blastema stages, similar to development, MEIS becomes limited to the proximal region while HOXA11 and then HOXA13 proteins are expressed in the distal zones of the mid-bud blastema (*30*) (*5*) (*29*) (*31*). These distal, HOXA13-expressing cells home to the regenerated hand even when transplanted into proximal regions showing they have acquired a stable identity (*29*). Over-expression of *Meis* and *Pbx1* in distal mid-bud blastema cells is sufficient to cause homing to the upper arm, demonstrating the functional importance of MEIS proteins in upper arm identity (*30*). Active MEIS-binding sites are found within the regulatory region of the gene encoding the cell surface protein PROD1 that is associated with upper arm homing (*32*).

Upper arm and hand blastemas display position-appropriate expression of homeoproteins after limb amputation (upper arm blastema expresses MEIS; hand blastema expresses HOXA13), but how the positional identity state is encoded in resting adult CT cells prior to regeneration has not been addressed. Here we performed chromatin profiling of CT cells in the three limb segments in axolotl and also during regeneration. We find that *Meis, Shox, HoxD9* and *HoxA13* homeoprotein genes show segment-specific accumulation of the repressive histone mark H3K27me3 that forebodes the upper arm versus hand expression of MEIS versus HOXA13. Typical upstream regulators of these homeoproteins do not show segment-specific profiles corroborating the developmentally quiescent state of the CT cells and their intrinsic properties upon transplantation. Regeneration involves transcriptional gene induction, and we observe the implementation of conserved developmental as well as non-conserved regeneration-specific enhancer elements harboring motifs for previously known and unknown conserved, regeneration-associated transcription factors. The regeneration-specific element at the *HoxA13* regulatory domain initiated its activity early in regeneration. This property together with the histone H3K27me3-low state of *HoxA13* specifically in the hand explains positional information and the segment-specific launching of the hand gene regulatory program.

## Results

### Homeodomain loci show histone modifications according to segment identity

To analyze limb segment-specific chromatin features, we used Fluorescence Activated Cell Sorting (FACS) to isolate CT cells from mature upper arm, lower arm, hand and head of the Prrx1:Cre-ERT; CAGGS:lp-STOP-lp-Cherry reporter axolotl (*33*). We performed Assay for Transposase Accessible Chromatin (ATAC)-Seq (3 replicates) and CUT&Tag for the histone marks H3K4me1, H3K4me3 and H3K27me3 (2 replicates) (Figure 1A, B, Supplementary Table S1) (*34*–*36*). Analysis of Accessible Chromatin Regions (ACRs) identified 1246 peaks that show reproducible differences between at least two segments listed above. Genomic feature annotation showed that 40% of peaks were found within 2 Kb of a gene body, with 60% found distally (Figure S1A). Hierarchical clustering of the differentially occurring ACRs (FDR<0.01, log2FC>1) after batch correction identified six distinguishable clusters including those with peaks strongest in the upper arm (cluster 1,2), versus those strong in the lower and hand (cluster 3,4) and those strongest in the hand (cluster 5,6) (Figure 1C, Supplementary Table S2). Interestingly, most differentially accessible peaks were associated with the hand and rather few with the upper arm. Within those peaks, active (H3K4me3), repressive (H3K27me3), and enhancer-associated (H3K4me1) histone marks were analyzed (Figure S1B, Supplementary Table S2). Comparison to the CUT&Tag profiles showed some correspondence of open chromatin peaks associated in a certain cell type with histone H3K4me3 or H3K4me1 marks. Conversely when that region is closed/lack of open peaks in a certain sample, the region is often associated with presence of H3K27me3 marks.

**Figure 1.**
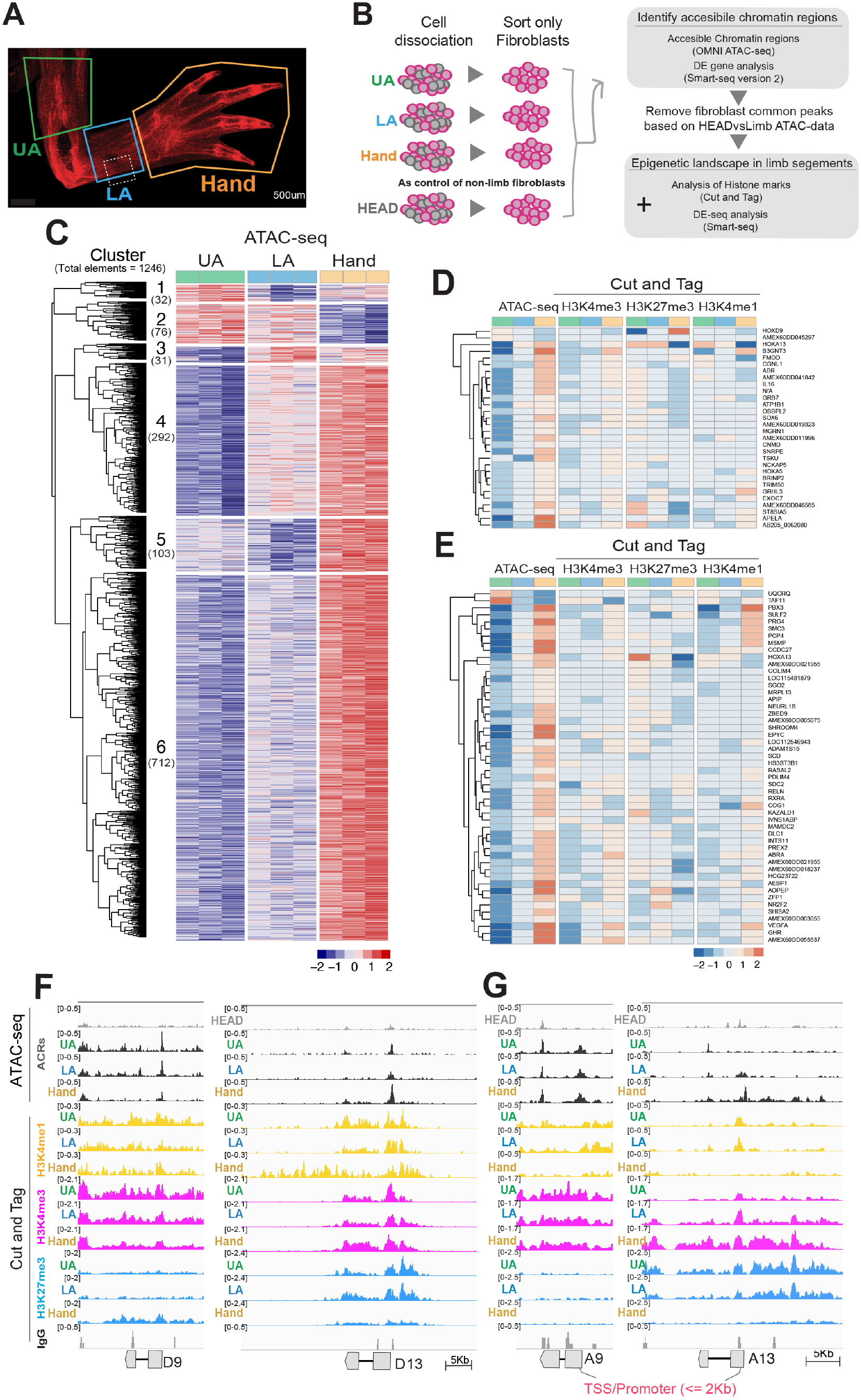
Axolotl limb CT cells show differential chromatin accessibility and histone H3K27me3 at *HoxD9* and *HoxA13* loci. (A) Wholemount image of a cleared limb of the *Prrx1:Cre-ERT; CAGGS:lp-STOP-lp-Cherry* reporter axolotl showing Cherry-expressing connective tissue (CT) cells. Each marked limb segment was used for cell isolation (Scale bar = 500 μm). (B) Schema of the experimental design. (C) Heatmap of segment-specific accessible chromatin peaks (1246) from axolotl mature upper arm (UA), lower arm (LA), and Hand CT cells. Six clusters were identified with hierarchical clustering using three replicates of ATAC-seq across segments. (D) Heatmap of promoter peaks in (A) (within 2 Kb from TSSs) that showed differential accessibility and their associated modified histone levels. *HoxD9* and *HoxA13* are among those showing strongest differential accessibility and differential histone H3K27me3. (E) Distal peaks showing differential accessibility and their candidate associated loci (top 50). (F) Profiles for accessible chromatin and histone modifications at the *HoxD9* and *HoxD13* loci show segment-specific differences. (G) Profiles for accessible chromatin and histone modifications at the *HoxA9* and *HoxA13. HoxA13* shows segment-specific differences. Accessible chromatin peaks were marked with dark gray, H3K4me1 with yamabuki-yellow, H3K4me3 with magenta, H3K27me3 with blue, and IgG control marked with light gray. The IgV tracks were shown where *HoxD9* and *HoxD13* (F) and *HoxA9* and *HoxA13* (G) are encoded.

To identify gene loci associated with segment-specific open chromatin, we identified open chromatin peaks within 2 kb of the annotated transcriptional start sites (TSSs) (Figure 1D, Supplementary Table S2). Given the different morphologies of the hand and upper limb skeleton, we expected to observe genes related to connective tissue differentiation in addition to potential positional information loci. Among the top 30 segment-associated promoter peaks, we found genes like *B3GNT3* a gene encoding an acetyl glucosamino transferase and *fibromodulin (Fmod*), a proteoglycan that modulates collagen fibril assembly (*37*). We noted that *HoxD9* and *HoxA13* homeoproteins were among the top three genes showing strongest segment differences in open chromatin (Figure 1D). Corresponding to their roles in development, the *HoxD9* promoter showed open chromatin in the upper arm segment, and inversely, higher repressive histone H3K27me3 levels across the entire gene in the hand (Figure 1F). Conversely, the *HoxA13* promoter showed open chromatin and histone H3K4me3 in the hand and higher histone H3K27me3 levels in upper and lower arm while *HoxA9* showed no noticeable differences (Figure 1D, G). Though conclusive assignment of distal peaks to genes is challenging, we made assignments using a probabilistic approach that takes into account the association between peaks and target expression within the same Topologically-Associating-Domain (TAD) (*38*) (and see Methods) (Figure 1E). An ACR between *HoxA11* and *HoxA13*, assigned to *HoxA13* was identified with open chromatin seen in the lower arm and hand, and conversely high histone H3K27me3 in the upper arm (Figure S1C).

We then asked whether there are gene loci showing differential histone H3K27me3 accumulation irrespective of open chromatin status at TSS, by comparing the magnitude of gene body-associated histone H3K27me3 between segments (Figure 2A, Supplementary Table S3). Multiple homeoprotein genes involved in limb development showed differential accumulation of histone H3K27me3, namely *HoxA11, HoxA10, HoxA13* (high H3K27me3 upper arm), and *Meis1, Meis2, Meis3, Shox, Shox2, HoxD8, HoxD9, HoxD10* and *HoxD4* (high H3K27me3 hand) (Figure 2A-C). Genes encoding cell surface proteins implicated in positional homing such as Tig1 (*9*) did not show detectable segment-specific differences.

**Figure 2.**
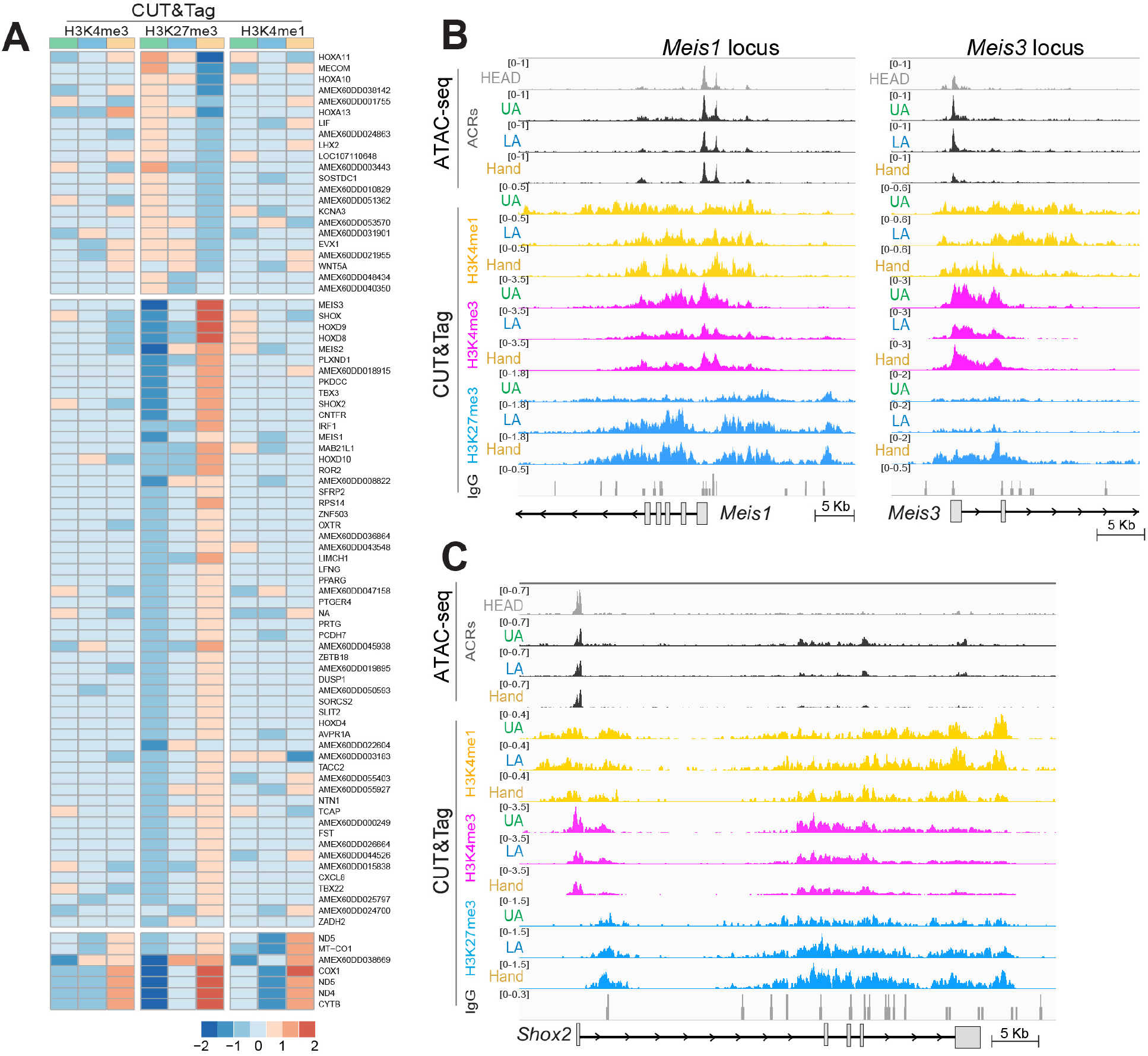
Differential histone H3K27me3 levels observed at multiple homeoprotein loci. (A) Heatmap of modified histone levels at genes where histone H3K27me3 levels integrated across gene bodies showed differential signal between segments. (B and C) Chromatin profiles of *Meis1* and *Meis3* (B) and *Shox2* (C) showing low levels of histone H3K27me3 in the UA, and higher levels in the Hand.

*Meis* and *HoxA13* mRNA and protein expression in histological sections is typically below detection levels in the mature limb and show a strong induction in the blastema (*6*, *29*). Nonetheless, given these segment-specific differences in repressive histone marks, we performed RNA-seq and examined transcriptional differences among the segments (Figure S2A, Supplementary Table 4). A global analysis showed that genes annotated as involved in chondrocyte and cellular differentiation showed the strongest representation in this dataset presumably reflecting the different skeletal morphologies and ratios of CT cell types in upper arm, versus lower arm versus hand regions (Figure S2B). As with the analysis of ACR at gene promoters, *Fmod* and *B3GNT3* were represented, as well as multiple collagen transcripts, *Col21A1, Col9A2, Col4A1 and Col7A1* (Figure S2A). *HoxA13, HoxD13, Meis1* and *Meis2* were identified as differentially expressed between hand and upper arm (Figure S2A). We additionally consulted a dataset in which a custom-designed microarray was probed with cDNA from CT cells from different segments. Again, cellular differentiation genes showed the strongest representation corresponding to the RNA-seq results (Figure S2C, D). Statistically significant differential signal was observed for *HoxA13, HoxD13, Meis1, Meis3, Shox* and *Shox2*,corroborating the segment-associated transcription of some of these limb homeoprotein genes in mature CT cells.

*Meis, Shox* and 5’ *HoxA* genes are expressed during limb development, and their upstream regulators have been characterized (*19*) (*28*) (*39*). Given the relatively low transcriptional status of these homeoprotein genes in mature CT cells, we wondered whether they might be uncoupled from the typical developmental regulatory inputs. In the developing and regenerating limb, RA is an upstream, proximalizing signal inducing *Meis* expression, counteracted by distalizing FGF, BMP and WNT3a signals (*12*, *13*) (*14*, *15*, *40*). We examined the RA target gene, *Rarβ* and observed no differential transcript abundance (Figure 3A, the genomic locus is unfortunately not present in the current version of the assembled genome). Furthermore, *Cyp26b1* which is expressed in the distal limb bud, and required for Meis repression (*18*), shows uniform histone marks (Figure 3B). For Fgf signaling, we examined *Spry4* and *Dusp6* and for Wnt signaling *Lef1* and *Axin2* as output and found no differential histone marks (Figure 3C, D). These results suggest that segment-specific features of these homeoprotein genes represent latent positional information.

**Figure 3.**
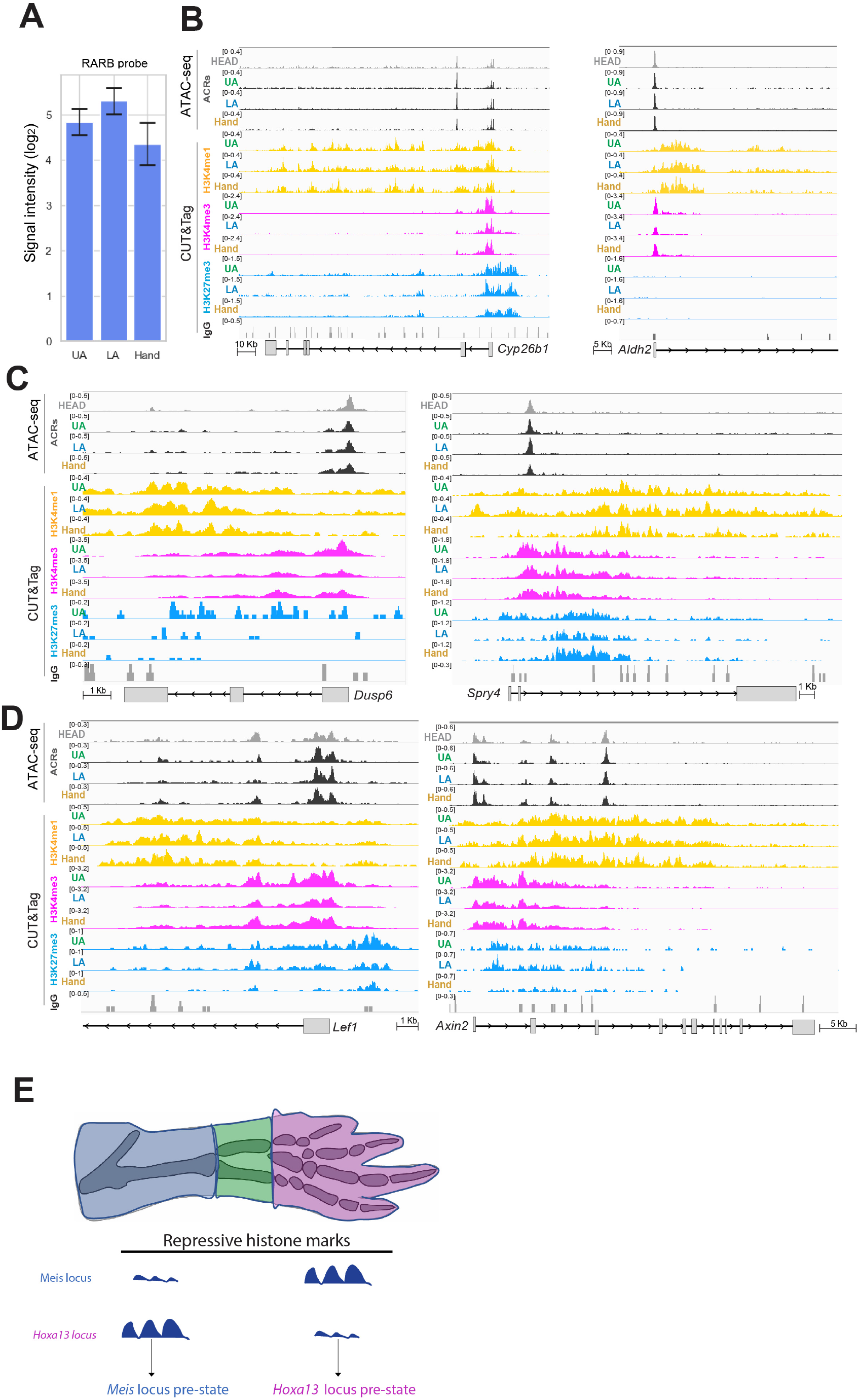
No evidence for segmental differences of typical developmental regulators of homeoprotein expression in mature limb CT cells. Examination in mature limb CT cells of target genes for several signaling pathways that normally regulate the proximo-distal spatial pattern of homeoprotein expression during development. (A) Bar chart of RARB signal intensity from limb CT cell microarray data (Log2 scale). (B - D) Chromatin profiles of target gene loci of several signaling pathways known to regulate homeoprotein expression during development. *Cyp26b1* and *Aldh2* as Retinoic Acid (RA) related genes (B), *Dusp6* and *Spry4* as FGF signaling downstream factors (C), and *Lef1* and *Axin2* as Wnt signaling downstream factors (D). *Lef1* is also a MEIS target. (E) Summary of differential histone H3K27me3 levels at *Meis* and *HoxA13* loci as pre-states of limb CT cells corresponding to future expression states in the UA versus Hand blastema.

Taken together, these results implicate selective, differential accumulation of repressive histones at homeoprotein genes yielding segment-specific transcriptional-ready states corresponding to “Positional Information” (*7*). We propose that the differential presence of repressive histone marks biases the launching of the transcriptional program after upper arm versus hand amputation (Figure 3E). After upper arm amputation, as *Meis* genes have low histone H3K27me3, they are easily activatable by amputation signals whereas *HoxA13* does not fire, as it is coated with histone H3K27me3. *HoxA13* would only become accessible upon re-entering a developmental growth progression. Conversely after amputation of the hand, since *HoxA13* has low occupancy of repressive histone marks, it is competent to respond to amputation signals, while *Meis* is not since it is occupied with H3K27me3 histones. Given the known feedback of HOXA13 protein as a repressor of upper limb-associated genes (*28*): (*27*), once the *HoxA13*-driven program initiates in a hand blastema, the upper arm program including *Meis* would not be initiated (Figure 3E).

### Identification of developmental versus regeneration-associated elements

We were therefore interested in chromatin features that change during regeneration. We performed chromatin profiling of a time course of upper arm regeneration in which we FACS-isolated CT descendants from the 5 (early bud), 9 (mid-bud) and 13 (palette, proximal and distal separated) day blastema (Figure 4A, B). Examination of *Meis1* showed maintenance of low histone H3K27me3 early in regeneration, and then acquisition of high histone H3K27me3 in the distal sample of the day 13 regenerate, corresponding to hand morphogenesis (Figure 4C). Examination of the *HoxA13* locus showed the maintenance of high histone H3K27me3 in the early blastema, corresponding to the lack of *HoxA13* expression, with a reduction of histone H3K27me3 in the distal sample of the day 13 regenerate (Figure 4D).

**Figure 4.**
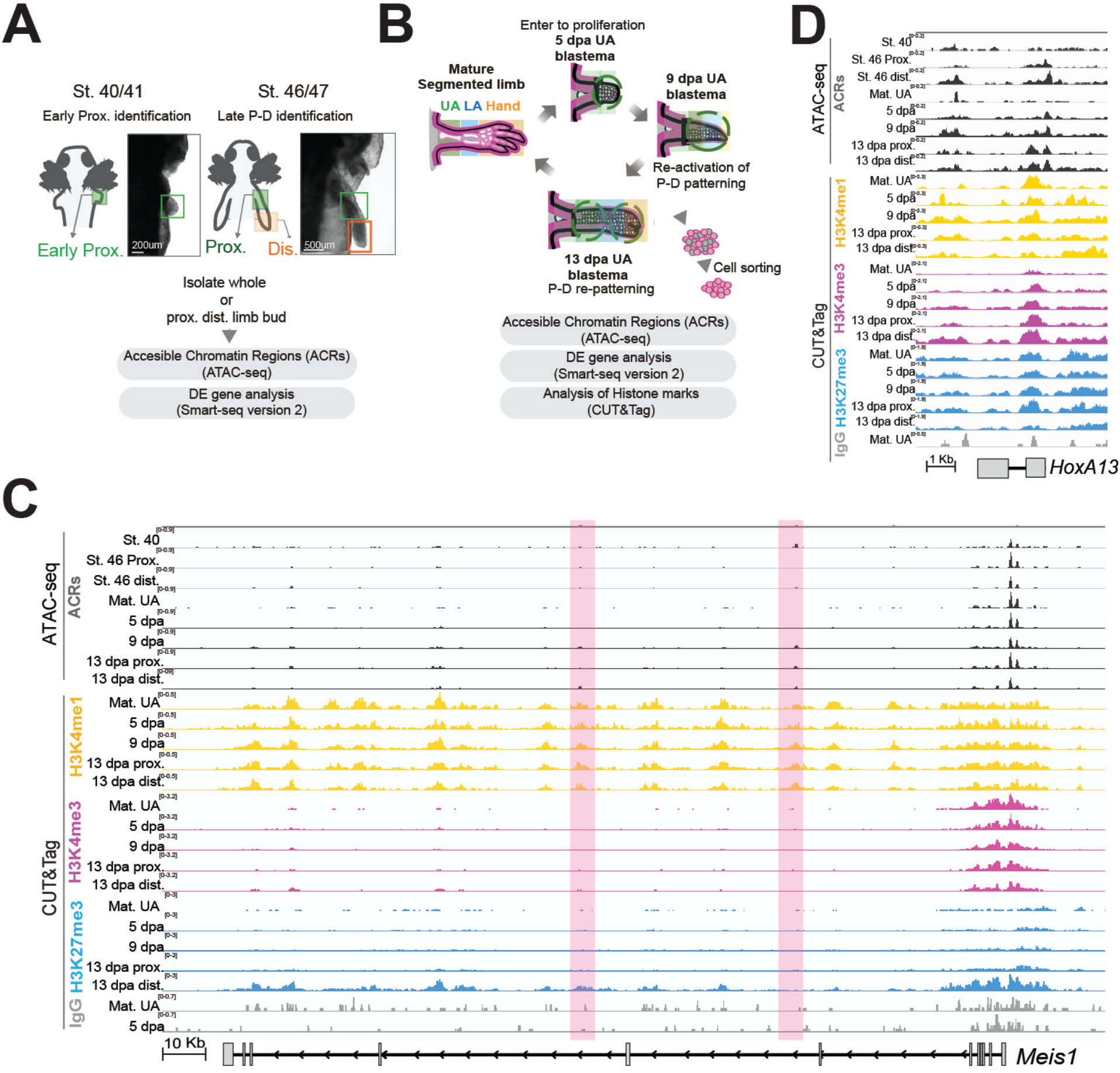
Chromatin profiling of CT descendants during upper arm regeneration. (A) Schema of profiling of embryonic limb buds at stage 40/41 and stage 46/47. ATAC-seq and Smart-seq v.2 libraries were generated from whole limb buds from stage 40/41, and separately from proximal or distal portions at stage 46/47. (B) Schema of profiling during a regeneration time course including ATAC-seq, Smart-seq v.2, and CUT&Tag. At days 5 and 9, cells were collected from whole UA blastemas. At 13 days the blastema was bisected into proximal and distal parts that were profiled separately. (C and D) Profiles from *Meis1* (C) and *HoxA13* (D) loci showing a change in levels of histoneH3K27me3 during regeneration in the distal 13 dpa blastema sample.

We then set out to explore genetic elements that are associated with gene expression during regeneration. To distinguish between genetic elements involved in redeployment of development versus initiating regeneration, we additionally performed ATAC-seq on two stages (stage 40 and 46) of the limb bud. Characterization of this dataset identified many (15763) regions whose chromatin state changed upon regeneration (Figure 5A, Supplementary Table 5). Clustering identified features common between the limb bud and the day 9 and 13 blastema, which we term Developmental Elements (DEs) (cluster 7, 8), while those that are only found in regeneration and not in the limb bud that we term Regeneration Elements (REs) (cluster 4) (Figure 5A). We explicitly examined the *Meis* and *HoxA13* regions. Among the DEs using VISTA analysis (*41*) we found 5 out of 19 Conserved Regulatory Elements (CREs) previously described for the mouse *HoxA* regulatory region including e2, e4, e5, e16 and e18 (Figure 5B, Supplementary Table 6). For example, axolotl e16 (axe16) showed accessible chromatin in the axolotl limb bud at Stage 40 but closed chromatin in mature, upper arm CT cells (Figure 5C, D). The region became open again in the day 9 blastema which corresponds to the developmental phase of regeneration (*33*). The axe16 region showed higher histone H3K27me3 in the proximal portion of the day 13 blastema, possibly reflecting downregulation of the activity during proximal morphogenesis.

**Figure 5.**
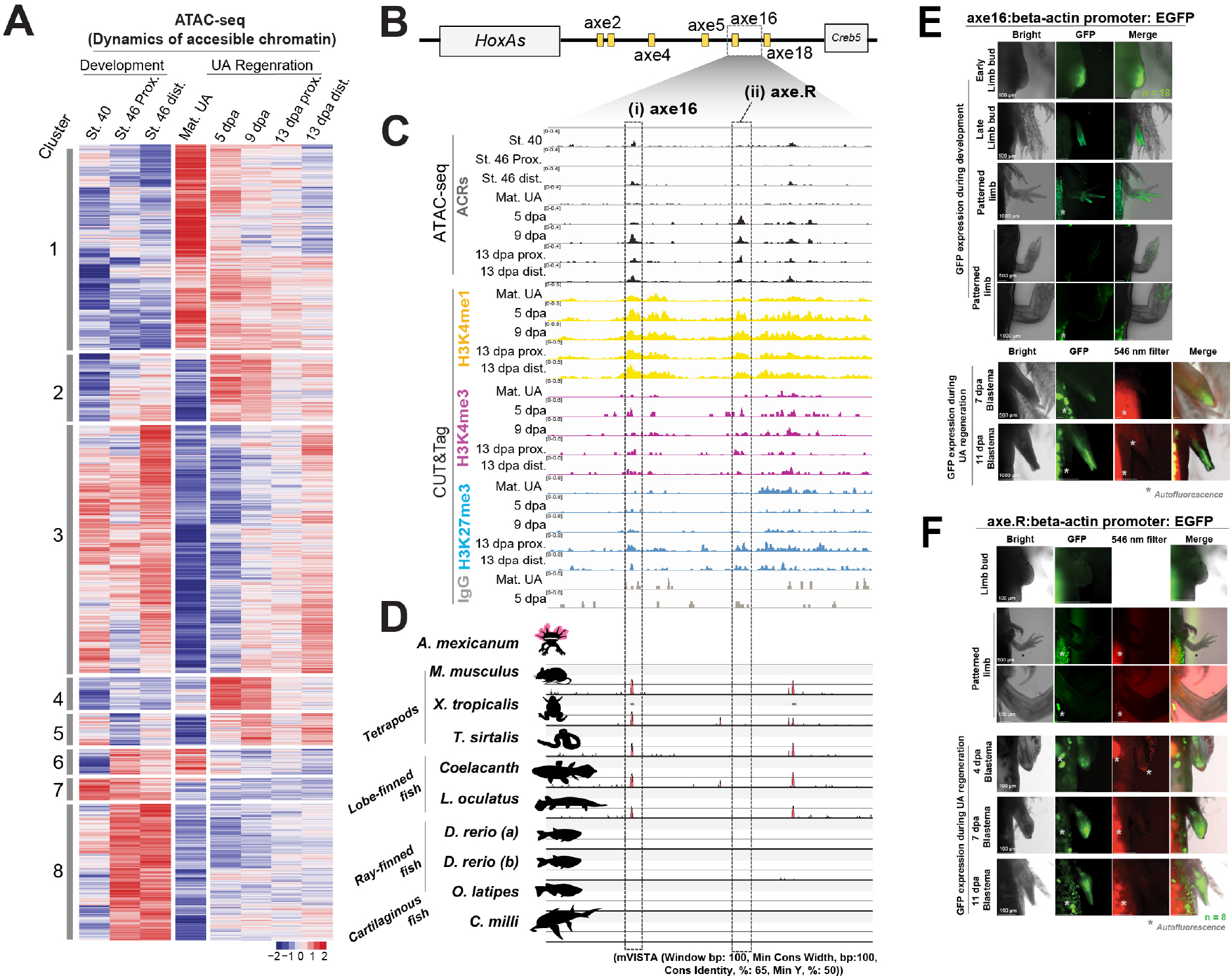
Identification of developmental and regeneration-specific elements that are activated upon upper arm regeneration. (A) Heatmap of dynamic ATAC-seq peaks (15763) across embryo stages, mature UA (Mat. UA), and regeneration timepoints of UA regeneration categorized in eight clusters. (B) Genomic organization of the *HoxA* cluster and putative regulatory region including conserved limb enhancers ((*1*)). Five axolotl genomic elements, termed *axe2, axe4, axe5, axe16*, and *axe18*, showed high conservation with previously identified mouse limb enhancers in mVISTA. (C) Chromatin profiles of developmental limb bud, Mat. UA, and regeneration timepoints in the genomic region surrounding *axe16*. Boxed areas denote the *axe16* (i), *axe.R* (ii) elements. *Axe16* showed ATAC-peaks in the limb bud and at later regeneration timepoints. *Axe.R* showed accessible chromatin peaks in early regeneration samples and not limb bud. (D) mVISTA alignment showing conservation rate at *axe16* and *axe.R* between axolotl and other vertebrate species. Axolotl genomic sequences were used as queries, and the corresponding genomic sequences from nine vertebrate species were used as subjects. *axe16* showed high conservation as putative regulatory elements of *HoxA* cluster genes between axolotl and three tetrapods and two lobe-finned fish, but not to ray-finned fish and cartilaginous fish (i). *Axe.R* did not show the sequence similarity between axolotl to any vertebrate species (ii). (E) *axe16* drives expression in the early to late limb bud, losing expression upon differentiation and was reactivated upon amputation in the regenerating blastema. (F) *axe.R* transgenic reporter animal drives GFP expression in the regeneration blastema but not the limb bud.

We looked within the regeneration-specific open chromatin datasets and found a RE (axe.R) in the distal *HoxA* regulatory region that showed no detectable sequence conservation with other tested vertebrates (Figure 5C, D). This element became accessible already by days 5 and 9 post-amputation (dpa) but was closed in the distal region at 13 dpa. Histone marks, H3K4me1, H3K4me3 showed increased in signal during regeneration, while H3K27me3 remained low during regeneration but appeared in the day 13 sample (Figure 5C).

To determine whether the axe16 and axe.R sequences functionally control developmental and regeneration-specific gene expression, we produced transgenic animals in which the genetic element was cloned next to a minimal actin promoter and the GFP sequence. Axe16:GFP reporter animals showed GFP signal in the mesenchymal limb bud from stage 40 to stage 52 whereupon GFP expression was downregulated after limb maturation/patterning (Figure 5E). Upon UA amputation, GFP expression reappeared in the blastema on 7 dpa and continued distally until 11 dpa (n = 18). In contrast, transgenic axe.R:GFP animals showed no GFP signal in the developing limb bud but upon UA limb amputation GFP expression was observed starting on 4 dpa (Figure 5F, n = 8). These results show that these elements function to control developmental and regenerative expression of target genes.

To probe the transcriptional program underlying the onset of regeneration, we performed motif enrichment analysis using MARA for peaks that showed differences in regeneration compared to mature (*42*) (Figure 6A). We also analyzed the RNA-seq dataset for transcripts encoding transcription factors (TFs) that arise in the regeneration time-courses (Figure 6B, Supplementary Table 7). The motif activity as well as transcript expression levels highlighted a set of 9 families of transcription factors, Runx, Bcl11, Bach, Fos/Jun, BATF3, MAFK, BNC and MAF_NFE2 as changing in the day 5 and 9 samples. Footprinting analysis of the ATAC-seq data sample confirmed potential time-dependent binding of, for example, Fos and Runx during regeneration (Figure 6C). Using this bulk ATAC-seq data together with previous scRNA-seq datasets (*33*) we could build transcriptional regulatory subnetworks active in mature versus regeneration timepoints (Figure S3A-E). This analysis showed strongest specific representation of TFs for *ETV6, NFIA* and *Twist2* in mature samples versus *Myc, Sall4, E2F3* and *HoxA11* in regenerating samples (Figure 6D, E).

**Figure 6.**
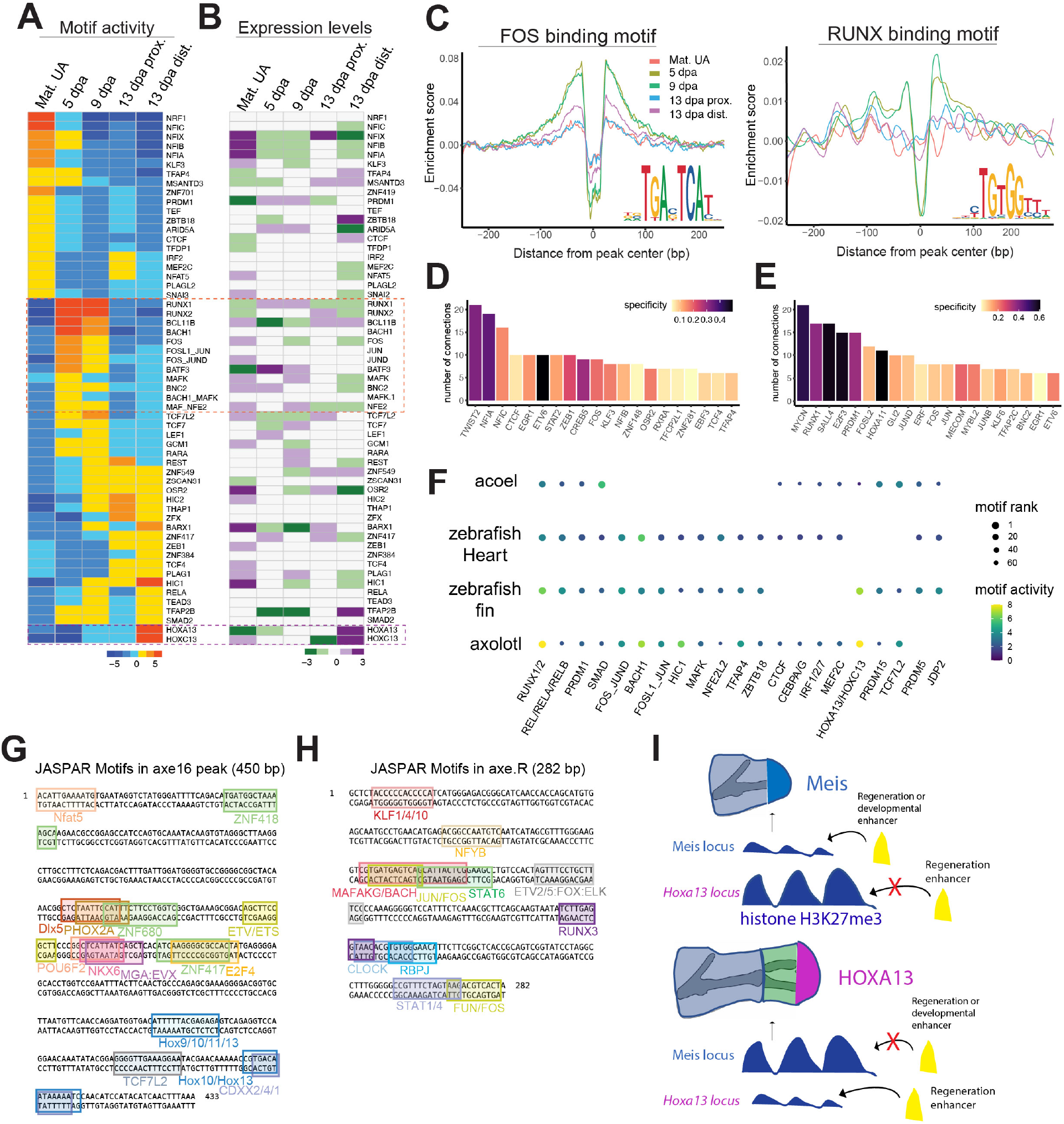
Transcription factor and motif analysis of regeneration regulatory network. (A) Motif activities (z-scores) inferred by MARA (see Methods) in Mat. UA and regeneration timepoints. (B) Associated TF expression from smart-seq v.2. (C) ATAC-seq footprints of FOS and RUNX motifs in Mat. UA and regeneration timepoints. (D-E) Connection numbers and specificities (ratio between Mat. UA-specific TF motifs and total) of top 20 TFs in Mat. UA-specific (D) and 5 dpa-specific (E) trimmed GRN. (F) Significant TF motifs inferred by MARA (see Methods) in the regeneration of axolotl limb, zebrafish fin, zebrafish heart and acoel. Motif ranks and activities were represented respectively by sizes and colors. (G and H) TF binding motifs from JASPAR analysis of axe16 (G) and axe.R (H), top 25 TF motifs are depicted. (I) Model for how segment-specific repressive histone pre-state modulates the ability to engage with enhancers. Amputation in the UA results in expression of *Meis*, but *HoxA13* is inaccessible due to high histone H3K27me3 levels. In contrast, in the Hand, *HoxA13* is accessible due to low histone H3K27me3 and can engage with early regeneration enhancers while *Meis* is occupied with repressive histones.

Regeneration has been profiled for chromatin accessibility and transcriptionally in other invertebrate and vertebrate species and we asked, to what extent these transcription factor motifs and transcription factors are present in other regeneration contexts. Strikingly, four out of the 22 regeneration-associated transcription factor motifs found in axolotl (*Runx, Rel, Prdm1* and *Smad*) were also found enriched in zebrafish fin and heart as well as acoel body regeneration (Figure 6F). These results point to a potentially metazoan-conserved regeneration launching program.

When we examined the presence of motifs in the *axe16* and *axe.R* elements, we observed motifs corresponding to their developmental versus regeneration status (Figure 6G, H). In particular, the *axe.R* element contained motifs for several of the transcription factors (*Jun/Fos, Mafk/Bach, Runx*) identified in the regeneration motif and transcription factor analysis (Figure 6G) meaning that a generic regeneration program could interface with poised positional information genes encoded by homeoprotein genes such as HoxA13.

We conclude that appropriate regeneration of the missing parts involves positional information in the form of repressive histone marks on homeoprotein genes corresponding to the role they played in development. In the hand, *HoxA13* with low histone H3K27me3 is accessible to REs like axe.R that induce an early onset of *HoxA13* expression launching the hand program. As HOXA13 suppresses the upper arm gene regulatory program, this early *HoxA13* induction in the hand prevents the upper arm program from launching in the hand (Figure 6I). During upper arm regeneration, regeneration enhancers including *axe.R* do not induce *HoxA13* expression in the upper arm since *HoxA13* is occupied by high levels of histone H3K27me3. *HoxA13* expression only initiates at the distal tip when histone levels are lower (Figure 4D) and when the developmental program has re-launched. These results implicate deposition of histone H3K27me3 as a key feature gating regeneration-specific and position-specific gene expression, coupled with the evolution of regulatory elements responsive to a regeneration transcriptional program that likely has been conserved from invertebrates to vertebrates.

## Supporting information

Supplemental Tables folder

## Acknowledgments

We thank the animal caretakers, Magdalena Blaschek, Tamara Torrecilla, Emina Silic and Viktoria Szilagyi for dedicated animal care. We are grateful to the Vienna Biocenter Facilities, NGS facility including Andreas Sommer, Alexander Vogt and Ido Tamir and the IMP/IMBA Molecular Biology Service including Harald Scheuch, Robert Heinan and Kristina Uzunova for their continuous support. We thank Maria Novatchkova, Thomas Burkhard, Siegfried Schloissnig, Diego Rodriguez Terrones, and Christiane Boeckel for analysis.

## Author contributions

Conceptualization: EMT, AK, DK, PM

Methodology: AK, JW, DK, PM, SN, WM, YTS, JF, EMT

Investigation: AK, JW, DK, PM, EMT

Visualization: AK, JW, EMT

Funding acquisition: AK, EMT

Project administration: EMT

Supervision: EMT

Writing – original draft: EMT, AK

Writing – review & editing: EMT, AK, JW, DK, PM, SN, WM, YTS, JF

## Data and materials availability

All data, code, and materials and methods used in the analysis are available in the Supplementary documents. Code is available at Github (https://github.com/labtanaka/positional_memory). Next Generation Sequencing raw data files have been deposited in GEO with number GSE217594:

SuperSeries: https://www.ncbi.nlm.nih.gov/geo/query/acc.cgi?acc=GSE217594

Linked SubSeries: https://www.ncbi.nlm.nih.gov/geo/query/acc.cgi?acc=GSE217591

https://www.ncbi.nlm.nih.gov/geo/query/acc.cgi?acc=GSE217592

https://www.ncbi.nlm.nih.gov/geo/query/acc.cgi?acc=GSE217593

## Supplementary Materials, Kawaguchi et al

### Materials and methods

#### Animal husbandry

d/d axolotls were maintained in individual aquaria and all animal breedings were undertaken by the IMP animal facility. All animal handling and surgical procedures were carried out in accordance with local ethics committee guidelines and as described previously (*1*). Animal experiments were performed as approved by the Magistrate of Vienna (License: GZ: 51072/2019/16). All animal surgeries and tissue amputations were carried out under anaesthesia in 0.03% benzocaine (SIGMA, E1501).

#### Generation of animal lines

Embryonic limb bud ATAC-seq and Smart-seq v.2 data were performed with the non-transgenic leucitic axolotl also known as *d/d* strain. ATAC-seq, RNA-seq, and CUT&Tag were performed using transgenic Prrx1:Cre-ER;CAGGs:lp-Cherry or Prrx1:Cre-ER;CAGGs:GFP-stop-lp-Cherry animals, in both lines mark limb fibroblast with Cherry due to the 4-hydroxytamoxifen treatment (*2*). The following publication describes the transgenesis and 4-OHT treatment in detail (*2*).

#### Generation of CAGGs:Lp-GFP-dead-Lp-Cherry animal

To generate, CAGGs:Lp-GFP-dead-Lp-Cherry animal, embryos of CAGGs:Lp-GFP-3xpolyA-Lp-Cherry animals were injected with a gRNA-Cas9 complex targeting GFP ORF (GGCCACAAGTTCAGCGTGTCCGG) as described previously (*3*). F0 animals that lacked GFP fluorescence were grown up to sexual maturity and they were mated to d/d animals to obtain F1 progeny. The F1 progeny was characterised using genotyping PCR to identify CAGGs:Lp-GFP-dead-Lp-Cherry animals. Sequencing of PCR product showed 9 nt deletion (TCAGCGTGT) at position 82 nt from ATG pertaining to EGFP. The F1 animals were further tested for their ability to convert to Cherry expression by injecting TAT-Cre protein injection. The established germ-line transmitted CAGGs:Lp-GFP-dead-Lp-Cherry line was used for mating with Prrx1:TFPnls-T2a-Cre-ERt to obtain cherry positive limb connective tissue cells.

#### Cell isolation and sorting for Chromatin profiling and Smart-seq ver. 2

The procedures of fibroblast isolation from limb tissue and blastema are described. In this study (*4*), we performed cell isolation with the following modifications: After cell dissociation with Liberase TM (SIGMA, 05401127001) treatment, cells were resuspended into 10%FCS/AMEM, and Fluorescence-activated cell sorting (FACS) was performed. Cell sorting with Aria III (AriaTMIII, BD biosciences) was carried out with optimal channels (488 nm and 567 nm), nozzle size was 85 μm and 0.8x FACSFlowTM was used at the IMP BioOptics facility. For CUT&Tag, mature ATAC-seq, and Smart-seq ver.2, mature upper arm and upper arm blastema cells (7.5 to 8cm animals from nose to tail) from 5, 9, and 13 dpa were isolated by FACS from the converted Prrx1:Cre-ER; CAGGs:lp-Cherry transgenic strain. For embryonic limb bud data with ATAC-seq and Smart-seq ver.2, the libraries were generated using cells isolated from white embryos. Stage 40/41 early embryonic limb bud cells were prepared from 40 whole embryonic limb buds. Late embryonic limb buds which were separated from proximal or distal parts were dissected from whole limb buds at Stage 46/47 embryos. The developmental stages of axolotl embryos were taken from (*5*).

#### Transgenic animal preparation

The transgenic constructs and animals were generated based on our recent paper (*6*). Axe16 and Axe.R were PCRed and cloned into the IS-bEGFP vector (*1*). The genomic sequence of axe16 was compared with vertebrates putative HoxA regulatory genomic sequences between *HoxA13* to *Creb5* (*7*) of 9 vertebrate species, *M. musculus, X. tropicalis, T. sirtalis, Coelacanth, L. oculatus, D. rerio, O. latipes*,and *C. milli* by mVISTA program (https://genome.lbl.gov/vista/mvista/submit.shtml). Axe16 was located at chromosome 2 (chr2p): 880953608 – 880953942 and Axe.R was located at chr2p: 880963799 – 880966329 (v.6.0-DD). The DNA fragments were amplified by Q5 High-Fidelity 2X master mix (NEB, M0492S) with the following primers: Axe16 with 5’-GCGgcggccgcACTTTAAAGCCCCAGATTAGGGTCG and 5’-GCGgcggccgcACGGTGTATGTCCTGGCCAGTC, Axe.R with: 5’-caccgcGTATTTGCCTGGGAGTAACCATGTCTC and 5’-cactagGGTTCCTGAGCTATTTGCAATTCTTAGGC. Then cloned into the IS-bEGFP vector. I-SceI-mediated transgenesis and Image acquisition were performed according to the previously described method (*1*) (*6*). In brief, DNA construct with I-SceI meganucrease was injected into single cell stage of white strain. F0 founders of axe16:EFGP transgenic animal were screened by EGFP fluorescence in early limb bud using a Zeiss Axio Zoom V16 stereo microscope. Since none of *axe.R:EGEP* F0 founder showed EGFP expression in the limb bud stages, around 24 animals were kept without fluorescence screening and raised in individual tanks. Their limbs were amputated at 2-3 cm from nose to tail, and EGFP expressions were observed in regeneration limb blastema.

#### ATAC-seq library preparation

ATAC-seq libraries were prepared based on the OMNI-ATAC seq method (*8*, *9*)with modifications as described below. Mature upper arm and upper arm blastemas (7.5 to 8 cm animals from nose to tail) from 5, 9, and 13 dpa were isolated by FACS from the converted Prrx1:Cre-ER;CAGGs:lp-Cherry transgenic strain as described. Embryonic ATAC-seq libraries were prepared using cells isolated from white embryos. Stage 40/41 early embryonic limb bud cells were prepared from 10 whole limb buds. All ATAC-seq libraries were performed from 1.5 to 2 ×10^4^ cells.

After FACS with Cherry positive populations or the isolated cells from embryonic limb bud, cell suspensions were spun down (250 G for 10 min at 4°C) and re-suspend in 500 μl Lysis buffer 1, mix gently and spun down (250 G for 10 min at 4°C). The pellets were resuspended in 100 μl Lysis buffer 2 and incubated on ice for 10 min. Lysis buffer 3 was added to the cells and mixed gently. Immediately, cells were spun down (250 G for 10 min at 4°C) and pellets were resuspended with 50 μl Tn5 Transposase reaction mixture (Transposase reaction: 5 μl of the 5x Transposase buffer (in-home production): 50 mM TAPS-NaOH (pH 8.5), 25 mM MgCl2, 50% DMF, 16.5 μl of PBS, 1 μl of 10% Tween-20 (0.1% f. c.), 1 μl of 1% Digitonin (0.01% f. c.), 2.5 μl of assembled in-house Tn5 (0.5 μg of assembled Tn5 was used for a reaction. Tn5 assembly with adapters was carried as described elsewhere (*10*). Transposition with Tn5 was carried out at 37°C for 1 hr with occasional agitation. The reaction was stopped by adding a 5 times volume of PB (QIAGEN) and vortexing for 30 seconds. Tn5 treated DNA was purified with MinElute PCR purification Kit (QIAGEN, # 28004) and eluted with 20 μl of EB buffer. 10 μl of purified DNA solution was used for library amplification, and final amplification cycles were defined with intermediate qPCR as described in the original ATAC-seq method (REF: Original ATAC). Nonetheless, all libraries were amplified within 11-12 total PCR cycles. Final libraries were amplified with NEBNext^®^ High-Fidelity 2XPCR Master Mix (NEB, M0541)). After PCR amplification, the libraries were purified with magnetic beads DNA isolation. The sequencing was performed using Hi-seq or Nova-seq PE125 or PE150 (the details are described in Supplementary Table 1) at VBCF-NGS facility.

#### CUT&Tag library preparation

CUT&Tag libraries were prepared using a protocol described by (*11*) (Bench top CUT&Tag V.2) with minor modifications as described below. To perform CUT&Tag libraries, 2 ×10^4^ cells of each mature segment (UA, LA and Hand) and regenerating UA blastema from 5, 9 and 13 dpa were corrected, and Cherry positive cells were sorted as described (*See in* Cell isolation and sorting). After sorting, cells were spun down and collected with 250 G for 10 min at 4°C and re-suspended into 100 μl (per sample) of 10% DMSO/AMEM with 10% FCS. Cells were frozen with Mr. Frosty Freezing Container (Thermo Fisher Scientific, 5100-0001) at −80°C until the library preparation started. All library preparation was started from frozen cells. Right before preparation started, cells were thawed on ice and spun down with 250 G for 10 min at 4°C. The pellets were re-suspended in 100 μl (per 2 ×10^4^cells) of pre-cold Nuclei Extraction with Protease inhibitor cocktail (Sigma, P8340-5ML) (NE with Pi) buffer and incubated on ice for 10 min with occasional agitation. Extracted nuclei were collected with 250 G for 10 min at 4°C and the nuclei pellets were resuspended in 100 μl (per 2 ×10^4^ cells) of Cold NE with Pi. Nuclei suspension was moved to PCR tube and mixed with 11 μl of Con-A beads (per 2 ×10^4^ cells) (Cell Signaling Technology, Ca. #93569) which were washed with buffer NE and activated following the manufacturer’s protocol. The nuclei resuspension and beads slurry were incubated for 10 min at room temperature. The tubes were placed on the magnetic stand until slurry clears and supernatant was removed. The pellet was washed twice with 50 μl of cold antibody150 (AB150) and gentle pipetting. The pellets were resuspended in 50 μl of cold AB150 and 0.75 μl of primary antibodies (H3K4me1/2: Abcam #ab8895) or 0.5 μl of primary antibodies (Normal rabbit IgG: Cell Signaling Technology #2729, H3K4me3: Abcam #ab8580, H3K27me3: Active Motif #39155) were added respectively and the tubes were incubated overnight at 4°C. The tubes were placed on the magnetic stand and washed twice with 100 μl of DIG150, and the beads were suspended in 50 μl of ice-cold DIG150. 0.75ul of secondary antibody (Epicyper: SKU #13-0047) were added and gently mixed and the tubes were incubated for 30 min at room temperature. The beads were placed on the magnetic stand and washed twice with ice-cold DIG300 and suspended in 50 μl of DIG300. 2.5 ug of pAG-Tn5 (Epicyper: SKU #15-1017) was added to the tube and gently mixed by pipetting. The tubes were incubated on the shaker for 1hr at room temperature. The tubes were returned to the magnetic stand and washed twice with 100 μl of DIG300. The beads were suspended in 50 μl of cold tagmentation buffer and incubated at 37°C for 1hr with occasional gentle agitation. The tubes were placed on the magnetic stand and the supernatant was discarded. The pellet was washed once with 50 μl of TAPS buffer by pipetting. The pellet was incubated with additional 5 μl of 10% SDS and incubated at 55°C for 1hr. The DNA was purified with QIAGEN MinElute PCR purification Kit (QIAGEN, # 28004) and eluted with 20 μl of 10mM Tris buffer (pH 7.4). 10 μl of purified DNA solution was used for library amplification. Final libraries were amplified with NEBNext^®^ High-Fidelity 2XPCR Master Mix (NEB, M0541)). 12 PCR cycles were used with all histone mark libraries, and 15 cycles were used with IgG libraries. Libraries were purified using magnetic beads DNA isolation. Sequencing was performed using the paired-end mode (the details are described in Supplementary Table 1) at VBCF-NGS facility.

#### Microarray Library design

A custom Agilent (www.agilent.com) microarray was designed using the axolotl transcriptome assembly v.25 that comprised several developmental stages (limb bud, tail bud, stage 10 and stage 19 (Bordzilovskaya, 1989)), regeneration stages (injured brain – 0, 3, 6, and 24 days post injury, 1 day injured spinal cord, 8 days tail blastema, 9 days and 15s day limb blastema), and mature tissues (heart, liver, lung, spinal cord, spleen, testes, brain of a metamorphosed axolotl). The RNA-seq data were assembled into contigs using Trinity (10.1038/nbt.1883). The contigs were annotated using a custom annotation pipeline.

For the annotated contigs that had a homolog in another organism, 3 microarray probes in the sense orientation and 1 in the antisense orientation were designed using the Agilent software. For those, without clear homologs, 3 sense and 3 antisense probes were designed, since the orientation of those contigs could not be determined reliably. Additionally, to ensure compatibility with the previous microarray studies in axolotl, we included 43736 designed earlier (*12*). Altogether, the microarray design comprised 415,996 different probes.

In this work, the microarray probes were mapped to the most recent genome and transcriptome assemblies, in order to assign them to the correct genes (see Segment-specific genes identified by microarray data of mature samples).

#### Tissue collection

Connective tissue of the limbs in the animals Prrx1:TFPnls-T2a-Cre-ERt, Caggs:LoxP-EGFP-LoxP-Cherry was converted from GFP-positive to Cherry-positive by 4-hydroxytamoxifen (4-OHT) treatment at the early limb bud stage as described previously (*2*) and tissue from the intact upper arm, lower arm and wrist areas was collected when the animals reached the size 4.5-5 cm from nose to cloaca (9-10 cm from nose to tail) and not sorted by sex.

Approximately 2 mm long sliver of tissue from the middle portion of the stylopod and zeugopod were collected for the upper arm (UA) and lower arm (LA) samples, and the portion of autopod containing two distal rows of carpals and metacarpals for the wrist sample. Tissue pieces from 6-8 animals from the same clutch were pooled to collect enough single cells.

Three biological replicates represent independently collected and processed samples from three separate animal batches.

#### Sample processing

Tissue was dissected into small pieced using forceps and dissociated to single cells for 30 minutes at room temperature in 1ml of dissociation solution containing 0.35 mg/ml Liberase TM (Roche# 05401119001) and 100 U/ml DNase1 (Roche# 04716728001) in 0.8x PBS. After approx. 30 minutes, tissue was further dissociated by pipetting, placed on ice and filtered through 70μm filter before FACS-sorting.

Cherry-positive cells were sorted directly into the Qiagen RLT buffer. Total RNA from 150,000 to 250,000 cells per sample was purified using Qiagen RNeasy mini kit according to the corresponding protocol.

Cy3-labeled cRNA probes were generated from 45 ng of purified total RNA using Agilent Low Input Quick Amp Gene Expression Labeling Kit. Custom designed 400K Agilent arrays (Name: Am400k_v2, Design ID:084163, Design Format: 2 x 400 K, Control Grid: IS-420288-2-V2_400Kby2_GX_EQC_20100210) were probed with 3.75 μg cRNA per array according to the Agilent protocol and scanned using the AgilentG3_GX_1Color Raw scan protocol. To normalize data between the arrays, we applied quantile normalization.

#### Smart-seq ver. 2 Library preparation

RNA isolation for embryonic limb bud, mature upper arm and upper arm blastema RNA-seq were performed from the same cell pools used for ATAC-seq samples. Around 1×10^4^ cells were spun down (450 G for 10 min at 4°C). Total RNA pools were isolated using an in-house RNA extraction kit. After the RNA isolation, RNA qualities of each sample were verified by the High-sensitive total RNA kit of Fragment analyzer (Agilent). Libraries were prepared with the Smart-seq v.2 protocol (VBCF-NGS facility) and sequenced with Nova-seq SR100 or PE100 platform by VBCF-NGS facility (the details are described in Supplementary Table 4).

#### Segment-specific genes identified by microarray data of mature samples

Microarray probes were first filtered if their sequences were mapped to axolotl transcriptomics (AmexT_v47) with more than 5 mismatches; and probes with detected expression levels significantly above the background (p-value > 0.05) were further selected. Then the background was subtracted from the probe intensities. Axolotl transcripts with less than 3 probes were further filtered. Data were normalized using the quantile normalization with the R package preprocessCore (v1.48.0). Pairwise comparisons of the mature samples (UA, LA and Hand) were done with limma (v3.42.2) and false discovery rates (FDRs) were calculated with Benjamini-Hochberge (BH) method. Segment-specific genes were identified if any pairwise comparison (LA vs. UA, Hand vs. UA or Hand vs. LA) was significant (FDR < 0.05 and log2 fold change > 1). The GO Term enrichment analysis of segment-specific genes was done with the R package clusterProfiler (v3.14.3).

#### Smart-seq2 data processing and analysis

Smart-seq2 processing pipeline was adapted for axolotl genome from nf-core/rnaseq (*13*). The raw reads were trimmed with trim_galore (v0.6.2) and mapped to axolotl genome (AmexG_v6 (*6*)) using hisat2 (v2.1.0). Read counts at gene levels were quantified by featureCounts in subread (v2.0.1). To identify segment-specific genes in mature samples, the read counts were normalized and pairwise comparisons were done with DESeq2 (v1.26.0) in the same manner as the microarray analysis. Genes with FDR <0.1 and log2FC >1 were considered significantly segment-specific. For regeneration-responsive genes, DESeq2 was also used to normalize the read counts and to compare all regeneration samples with the mature UA, while the batch effect was taken into account in the design matrix. The regeneration-responsive genes were then defined as the differentially expressed (DE) genes with FDR <0.05 and log2FC > 1 in at least one comparison. The regeneration libraries of the batch-corrected gene expression were obtained with Combat in sva package (v3.34.0).

#### ATAC-seq data analysis and segment-specific chromain accessible regions in axolotl mature samples

The pair-ended ATAC-seq reads were trimmed with cutadapt (v1.18) and aligned to axolotl genome (AmexG_v6) with bowtie2 (v2.3.5.1). Multimappers and duplicated reads in the aligned bam files were discarded respectively with samtools (v1.10) and picard (v2.20.6). ATAC-seq narrow peaks were called by macs2 (v2.2.5) with default parameters. The processing workflow was wrapped with nextflow (*14*). For the purpose of reproducibility, peaks were retained only if they were called in at least two biological replicates with p-value < 106; they were merged into a list of peak consensuses for all samples. Read counts were quantified within those peaks with feature counts from subread (v2.0.1). After low-signal peaks were filtered with less than 50 read counts, the read counts within peaks across all samples were normalized using the scaling factors from DESeq2 (v1.26.0). These scaling factors were also used to make bigwig files with deeptools (v3.1.2) for the data visualization. To mitigate the batch effect in the peak signals in the mature samples, we applied Combat in sva package (v3.34.0). Genomic features of ATAC-seq peaks were annotated with ChIPseeker (v1.22.1) using annotated 23585 fibroblast-expressing genes in axolotl limb based on our microarray and smart-seq2 data. The pairwise comparisons were performed with edgeR (v3.28.1) using the batch-corrected peak signal. The segment-specific ATAC-seq peaks were defined if any pairwise comparison (LA vs. UA, Hand vs. UA, or Hand vs. LA) was significant (FDR < 0.05 and log2FC > 1). The obtained peak list were further filtered if they were overlapped by mature head samples. Finally, 1246 segment-specific peaks were found and grouped into six clusters with the hierarchical clustering.

#### Dynamic chromatin accessibility in axolotl limb regeneration

To determine the chromatin accessibility landscape in axolotl limb development and regeneration, we first applied Combat in sva package (v3.34.0) to the ATAC peak signals of regeneration and development to removal the batch effect and then we compared all samples with mUA using edgeR (v3.28.1). Dynamic peaks or CREs were identified with FDR < 0.01 and log2FC >1 in at least one of those comparisons. To group those peaks across time points with DPGP clustering approach (*15*), the biological replicates were first averaged and down-sampled to 10,000 peaks for the sake of algorithm efficiency. With given clusters from DPGP, we calculated the correlations between profiles of non-selected peaks and average profiles of each cluster and assigned peaks to the clusters with the highest correlation and the correlation higher than 0.6. To reduce the complexity, clusters with the small number of peaks were manually merged to the most similar ones. Finally, 15763 dynamic CREs were grouped into eight clusters.

Footprint analysis was performed with TOBIAS (v0.13.3) (*16*) using the vertebrate motifs from the CORE collection of JASPAR2022 (*17*).

#### CUT&Tag data processing and analysis

The raw data of CUT&Tag were processed with the same next-flow pipeline as ATAC-seq; the peak consensus for all three histone marks (H3K4me1, H3K4me3, and H3K27me3) were also identified in a similar manner as ATAC-seq (see above). Notably, all previously ATAC-seq peaks were included in the histone mark peak consensus. Read counts within histone peaks were quantified with feature counts from subread (v2.0.1) and were normalized using the scaling factors from DESeq2 (v1.26.0). The same scaling factors were also used to make bigwig files with deeptools (v3.1.2) for visualization except the IgG control samples, which were normalized with library size in deeptools. The batch effects in the histone peak signals were corrected with Combat in sva package (v3.34.0) respectively for each histone mark. In addition, segment-specific and regeneration-responsive histone marks were also determined in the similar way as ATAC-seq data (FDR < 0.05 and log2FC > 1).

For the gene-based histone analysis in the mature samples, annotated genes in axolotl genome (AmexT_v47) and their 5kb upstream were considered. Read counting, normalization, batch correction and pairwise comparisons were done in the same way as before. To account for different gene length, the histone mark levels were further normalized by length (RPKM (*18*)) and lowly expressed genes were filtered with log2rpkm <= 0.6.

#### Motif Activity Response Analysis (MARA) for regeneration in axolotl and across species

To infer TF motif activities involved in axolotl limb regeneration, we adapted MARA (Motif Activity Response Analysis) described in (*19*). Specifically, we modeled the ATAC temporal signals within the regeneration peaks (in log2 scale) as a linear combination of TF binding sites and motif activities (*20*):

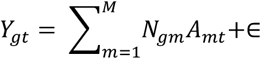

where *Y_gt_* is the ATAC signals (in log2 scale) in genomic regions g and at regeneration time point t; *N_gm_* is the number of TF binding sites in the region g for TF motif m scanned by FIMO using JASPAR2022 CORE motifs; *A_mt_* is the activity of motif m at time t; *ϵ* is the gaussian noise. To control for over-fitting and redundancy of motifs, we employed Bayesian ridge penalty in the linear regression (implemented in (*21*)). The resulting motifs were further filtered if corresponding TFs were not expressed in the Smartseq2 data. For enrichment scores of axolotl regeneration-specific ATAC peaks (cluster 4 in Fig2C), fisher exact p-values was calculated. For the crossspecies motif analysis, ATAC-seq and RNA-seq data of zebrafish fin (*22*), zebrafish heart (*23*) and acoel (*24*) were processed and analyzed in the same way as axolotl; and MARA was performed to infer temporal motif activities.

#### A statistical method for enhancer-to-gene assignment accounting for the correlations within Topologically Associating Domains (TADs)

To improve the distal enhancers-to-gene assignment, we implemented the statistical approach in (*25*). Specifically, all ~50,000 ATAC-seq peaks were first annotated with ChIPseeker (v1.22.1) and those annotated as ‘Distal Intergenic’ and ‘Intron’ were selected as potential enhancer candidates and further filtered based on enhancer mark H3K4me1 levels (logRPKM > 1). To determine the target of each enhancer, only genes within the same axolotl TADs (resolution of 100kb identified in (*6*)) were considered. In addition, each chromatin feature (ATAC, H3K4me3, H3K27me3 and H3K4me1) was used to calculate the correlations with the candidate expression measured by Smart-seq2. The ATAC-seq signals were selected because of the highest correlations with the gene expression. Finally, only the significant (p-value <0.05 with Student’s t-distribution) candidate with the best correlation was assigned to the enhancer, otherwise no target was assigned.

#### Infer transcriptional regulatory network (TRN) of Axolotl limb regeneration by combining bulk ATAC-seq and single cell RNA-seq

To infer the gene regulatory network of axolotl limb regeneration, we integrate bulk ATAC-seq data and the scRNA-seq data (*2*). The main steps of our approach were as follows:

1. TFs relevant to axolotl limb regeneration were collected first. We started with the full TF list from (*26*) and kept TFs that were detected in scRNA-seq data (blastema data points 0, 3, 5, 8, 11, 18dpa); TFs were further selected if they were previously identified by MARA analysis, or as segment-specific genes, or as regeneration-response genes, or annotated as limb development in GO Term.
2. To maximise the number of TFs with available motifs, the JASPAR2022 CORE and UNVALIDED motifs were collected; and motifs in the CORE were selected if TFs had motifs from both collections. Promoters (−2000-+300bp from the annotated transcriptional start site (TSS)) and enhancers peaks from ATAC-seq data that were assigned to axolotl limb expressed genes were scanned for TF motifs using FIMO.
3. By associating motifs found by FIMO to previously selected TFs, gene targets and potential regulators (TFs) matrix was obtained and served as prior network. To reduce the complexity, we were solely interested in the transcriptional regulatory networks (TRN) and thus only TFs themselves among gene targets were retained.
4. For each target, to select the important regulators, source code of GENIE3 was modified to adapt the prior knowledge of potential regulators. The random forest approach was used with 1000 trees. The resulting edges were further filtered with importance threshold of 0.01. To visualise the inferred GRN, the embedding UMAP (R package uwot, v0.1.11) was computed based on the principal components of the scRNA-seq data of TFs. Network modules were done using igraph (v1.3.1). The regeneration-time specific GRNs were pruned based on the sample-specific chromatin accessibility in the similar manner as (*27*).

#### Data and materials availability

Code is available at Github (https://github.com/labtanaka/positional_memory). Fastq files have been deposited in GEO with number GSE217594: SuperSeries: https://www.ncbi.nlm.nih.gov/geo/query/acc.cgi?acc=GSE217594

Linked SubSeries: https://www.ncbi.nlm.nih.gov/geo/query/acc.cgi?acc=GSE217591

https://www.ncbi.nlm.nih.gov/geo/query/acc.cgi?acc=GSE217592

https://www.ncbi.nlm.nih.gov/geo/query/acc.cgi?acc=GSE217593

All other data are in the main paper or supplementary materials.

**Figure S1.**
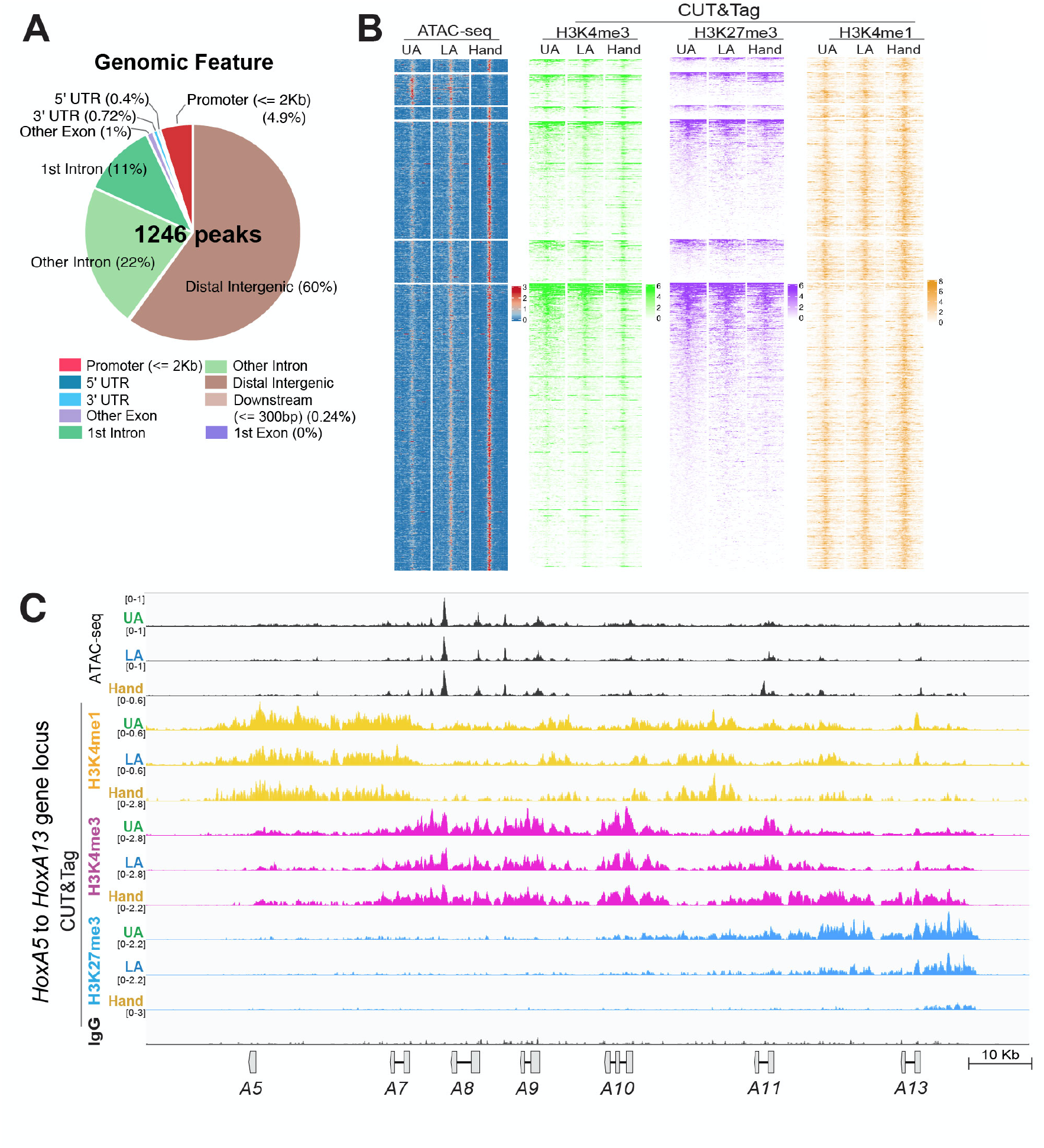
Genomic and chromatin features of UA, LA, Hand CT cells. (A) Pie chart of genomic annotation of 1246 segment-specific ATAC-seq peaks in Figure 1C. (B) Genomic coverage heatmap of segment-specific ATAC-seq peaks (1246) and associated histone marks (H3K4me3, H3K27me3, and H3K4me1) in axolotl mature UA, LA and Hand. Six clusters were identified with hierarchical clustering using three replicates of ATAC-seq across segments, as in Figure 1C. (C) IgV tracks between different segments with ATAC-seq and histone modification profiling of HoxA gene cluster.

**Figure S2.**
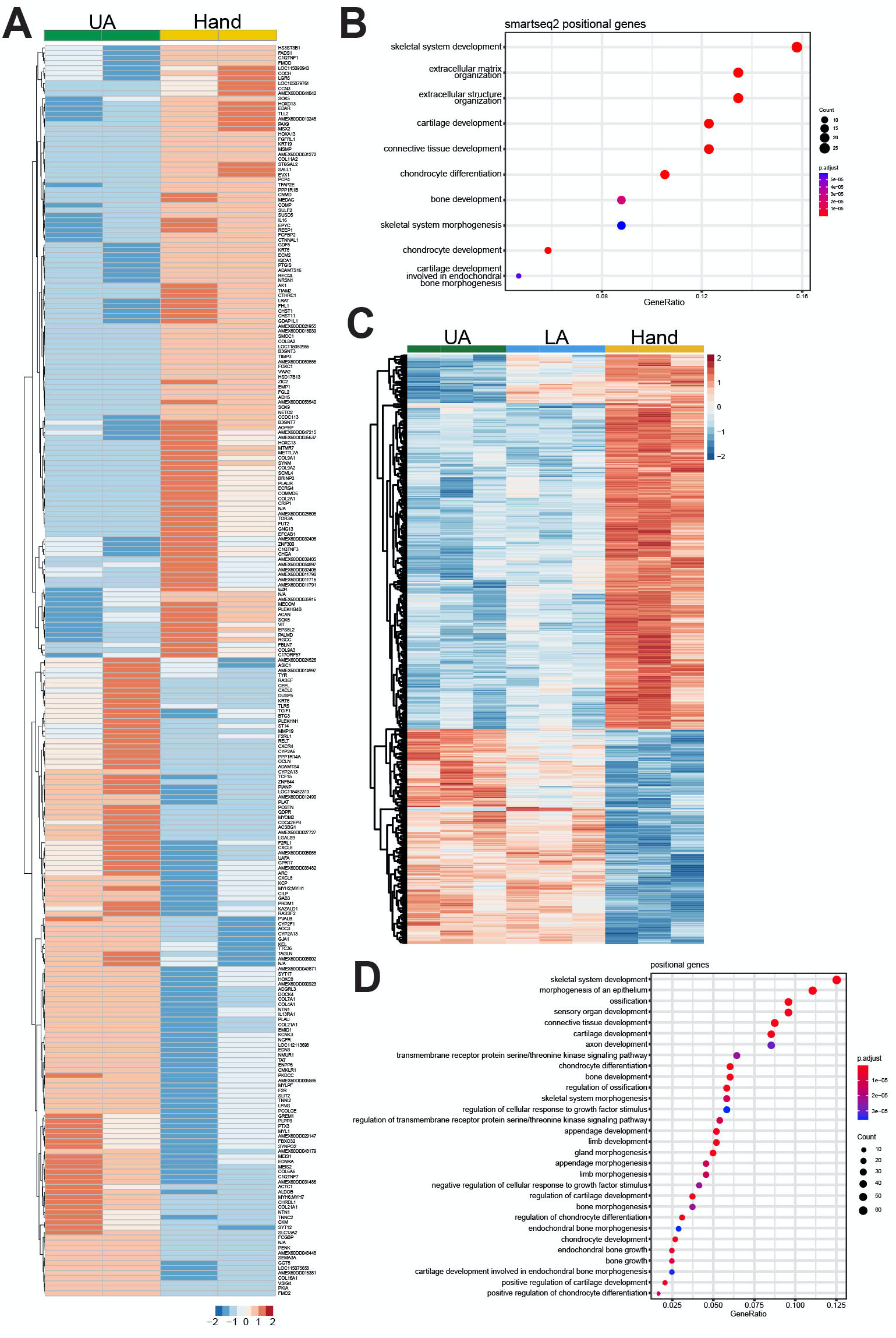
Transcriptional analysis of UA versus Hand CT cells. (A) Heatmap of differentially expressed (DE) genes between mature UA and Hand (FDR <0.1 and log2FC > 1) by Smartseq2 data. (B) GO Term enrichment of DE genes in panel A. (C) Heatmap of differentially expressed (DE) genes between mature UA, LA, and Hand (FDR <0.05 and log2FC > 1) in an independent microarray experiment. (D) GO Term enrichment of DE genes in the panel C.

**Figure S3.**
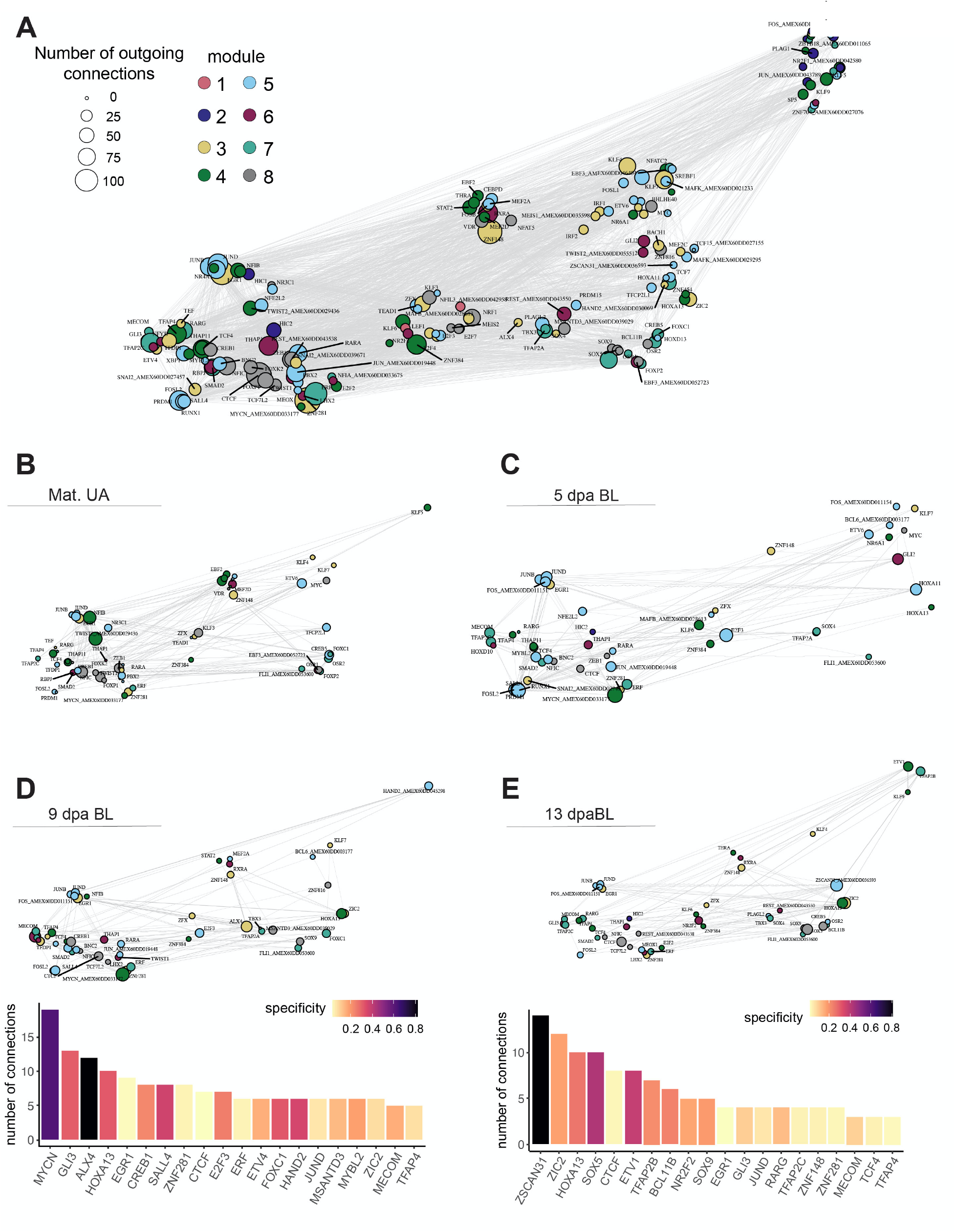
Gene regulatory network of limb regeneration. (A) UMAP embedding of inferred gene modules based on the TF-to-gene associated from bulk ATAC-seq and coexpression in scRNA-seq data. Gene modules are color-encoded and node sizes represent the number of outgoing connections of TF. (B-C) UMAP embedding of trimmed GRN based on mat. UA (or 5dpa-specific) chromatin open TF regulatory regions; gene modules are color-encoded and node sizes represent the number of TF connections. (D-E) Upper panels: UMAP embedding of trimmed GRN based on 9 dpa-specific (or 13 dpa-specific) chromatin open TF regulatory regions; gene modules are color-encoded and node sizes represent the number of TF connections; lower panels: connection numbers and specificities of top 20 TFs in 9dpa-specific (or 13 dpa-specific) trimmed GRN.

**Table S1: Summary of ATAC-seq and CUT&Tag information from mature limb segments.**

Sheet1: NGS detailed information of ATAC-seq libraries from mature limb segments.

Sheet2: NGS detailed information of CUT&Tag libraries from mature limb segments.

**Table S2: Differential chromatin accessibility in axolotl limb CT cells.**

Sheet1: Segment-specific accessible chromatin peaks (1246) identified from axolotl UA, LA and Hand (Figure 1C) and peak annotation.

Sheet2: ATAC-seq and histone mark levels of top 30 segment-specific peaks annotated as promoter peaks (Figure 1D).

Sheet3: ATAC-seq and histone mark levels of top 50 segment-specific peaks annotated as enhancer peaks (Figure 1E).

**Table S3: Analysis of differential histone marks from CUT&Tag between Mature limb segments.**

Sheet1: 86 gene-centric differential histone marks across axolotl segments (Fig 2A)

**Table S4: Segment-specific genes in axolotl limb CT cells identified in Smart-seq ver. 2 and microarray data.**

Sheet1: Smart-seq ver. 2 sequencing data set

Sheet2: Gene expression (log2 scale with DESeq2 normalization) of 247 segment-specific genes identified in Smart-seq ver. 2 between UA and Hand with FDR<0.1and log2FC >1.

Sheet3: Gene expression of 717 segment-specific genes identified in microarray data with FDR<0.05 and log2FC >1.

**Table S5: Clustering of dynamic chromatin changes across developmental stages and regeneration time points.**

Sheet1: 15763 dynamic ATAC-seq peaks across embryo stages, mature UA and regeneration timepoints (Fig 5A).

**Table S6: Genomic coordinates of conserved regulatory elements at HoxA limb enhancers.**

Sheet1: Genomic coordinates of conserved HoxA regulatory elements that identified in this study.

**Table S7: 4843 regeneration-responsive genes in axolotl CT cells across regeneration time points from Smart-seq ver. 2.**

