## Supplemental Tables folder for "Chromatin states at homeoprotein loci distinguish axolotl limb segments prior to regeneration": Supp_table6 copy.pdf

| No. of Chr. | Start | End | Name of CNE from mouse* |
|---|---|---|---|
| chr2p | 875846298 | 875847543 | axe2** |
| chr2p | 875800180 | 875801429 | axe2** |
| chr2p | 876688104 | 876688398 | axe4 |
| chr2p | 877765382 | 877765724 | axe5 |
| chr2p | 880953608 | 880953942 | axe16 |
| chr2p | 881033675 | 881033889 | axe18 |
| chr2p | 880963799 | 880966329 | ax.R |

* Conserved elements were exported from mouse paper (Berlivet S. et al., PLOS GENETICs (201

** There are two genomic loci in Axolotl genome 6DD

3))
